# A multi-species repository of social networks

**DOI:** 10.1101/464271

**Authors:** Pratha Sah, José David Méndez, Shweta Bansal

## Abstract

Social network analysis is an invaluable tool to understand the patterns, evolution, and consequences of sociality. Comparative studies over the spectrum of sociality across taxonomic groups are particularly valuable. Such studies however require quantitative information on social interactions across multiple species which is not easily available. We introduce the Animal Social Network Repository (ASNR) as the first multi-taxonomic repository that collates more than 650 social networks from 47 species, including those of mammals, reptiles, fish, birds, and insects. The repository was created by consolidating social network datasets from the literature on wild and captive animals into a consistent and easy-to-use network data format. The repository is archived at https://bansallab.github.io/asnr/. ASNR has tremendous research potential, including testing hypotheses in the fields of animal ecology, social behavior, epidemiology and evolutionary biology.

## Background & Summary

Network analysis is rapidly becoming a central approach in several basic and applied research areas of ecology and evolutionary biology, including behavioral ecology, epidemiology, spatial ecology, and social evolution [14]. Recently, researchers have demonstrated the utility of network analysis in explaining the transmission of social information in animal groups [13, 4], evolution of cultural behavior [17], epidemiological consequences of group substructure [19], mechanisms of infectious disease transmission in animal groups [22, 19, 8], and emergence of collective behavior [5]. Network analysis leverages detailed social interaction and movement data and allows for the incorporation of heterogeneity at the individual scale to explain population level processes, as well as the ability to objectively quantify the organization of social interactions and dynamics of group behavior.

Recent advances in computational power and technological tools, such as proximity loggers and radio-frequency identification, have facilitated the collection of network data [15]. And with the open science movement gaining steady momentum [11], a culture of making research data and experimental methods publicly available and transparent has unleashed valuable social interaction data for use by all researchers. For the first time, there is thus an opportunity to carry out comparative network studies across multiple species to identify general patterns and generate broad principles [19]. However, such studies are currently challenging due to differences in data collection methods, a lack of standardized formats for published data, and the absence of a centralized data repository [20]. While such repositories exist for human interaction data [2, 3, 16, 1], there is a gap for social network datasets across multiple taxonomic groups.

Here, we introduce the Animal Social Network Repository (ASNR), which fills this gap by providing access to networks of social interactions across multiple taxonomic groups organized in a consistent network file-format. The repository provides opportunities for: (a) field biologists to generate preliminary hypotheses and plan for data collection (including the resolution, duration and quality) required to test their hypotheses; (b) empiricists to evaluate the effects of data collection methods on observed network properties and characteristics;(c) behavioral ecologists to compare social structures within and across broad taxonomic groups; (d) network scientists to analyze the patterns and function of dynamic networks; (e) evolutionary biologists to understand the drivers for the emergence of disparate network structure across different species; and (f) and disease ecologists to understand the eco-epidemiological implications of the evolved network structure. The repository thus has tremendous research potential in the fields of ecology, epidemiology, evolution, behavior, and beyond.

## Methods

### Selection methodology

The data repository was collated by reviewing published literature and popular data repositories, including the Dryad Digital Repository, Harvard Dataverse and figshare, for social network datasets associated with peer-reviewed publications. We used terms such as “social network”, “social structure”, “interaction network”, “animal networks”, “network behavior”, among others, to perform our electronic search. Additional network datasets were acquired by directly contacting the authors of published studies without open data. Only studies on non-human species were included, and studies reporting non-interaction networks (i.e. biological networks or food-web networks) were excluded. Studies that did not include enough information for networks to be re-created were also excluded. By reviewing the quality of the remaining published datasets (see Technical Validation), a total of 666 social networks spanning 47 animal species and 18 taxonomic orders were selected for the data repository.

## Data Records

The social network files from this study are available through the ASNR website (https://bansallab.github.io/asnr) and Harvard Dataverse (https://doi.org/10.7910/DVN/N5YHLL). The ASNR website serves as a dynamic platform, where new network datasets can be curated and added. For easy access to the datasets, the repository is organized into sections each representing a unique taxonomic group. Each section further consists of a set of social networks which were collected together with the same sampling method. The datasets are uniquely identified with the animal species first, followed by the interaction type and ending with the edge weight criteria (weighted or unweighted). In cases where multiple networks are available within each dataset, each social network is assigned a unique identifier. A readme file is also included with each dataset that summarizes structural features of the networks and provides information on the original source.

Each network dataset is provided in the `GraphML` format [7]. `GraphML` is a flexible and convenient XML format for storing network information. It supports unweighted, weighted, undirected and directed networks and allows for the definition of node and edge attributes.

## Technical Validation

Our validation process consisted of data-type and constraint validation, structural validation, and cross-reference and ecological validation. All data collection and validation steps were carried out by two co-authors (PS and JM).

### Data-type and constraint validation

The first step involved quality checks to ensure that the original data contained enough information to enable reconstruction of social network(s). All datasets were acquired in electronic format in one of the following four network data structures: edgelists, adjacency matrices, adjacency lists or group membership dataframes. All data was classified into nodes, edges or attribute data. All node ids were verified to be of the same type (e.g. integer or string). All edges were verified to be between nodes in the node list, or were added as nodes to the node list. All attribute data was verified to correspond to an existing node or edge.

### Structural validation

We next validated the structural integrity of the network described in the original data-source by removing all edges that connected any node to itself (i.e. self loops). Any duplicate edges were also removed. Individuals with no edges (i.e. isolated nodes) were not removed from the network.

The ASNR currently only contains static networks. Thus, multiple interactions reported between the same node pair at different time-points were replaced with weighted edges, with weights representing the interaction frequency.

### Cross-reference and ecological validation

For detecting errors in the data mining and `GraphML` conversion process, we calculated network summary statistics (e.g. number of nodes, number of edges, clustering coefficient) for each network and cross-checked them against the network description in the original publication. The structures of each converted network file were also cross-checked to ensure consistency within the ecological context of data collection. For example, networks of the same group of individuals of a species that were collected over mating vs. non-mating season are expected to differ in terms of their network densities.

## Data Characterization

In the sections below, we characterize the phylogenetic and geographical distribution, data collection methodology, and structural similarity of the networks included in the repository.

### Phylogenetic and geographic distribution

The phylogenetic distribution of the taxonomic groups currently included in the repository is shown in Figure 1. While mammals are the most studied taxa, social networks from other taxa including reptiles, birds, insects, and fish also exist.

**Figure 1:**
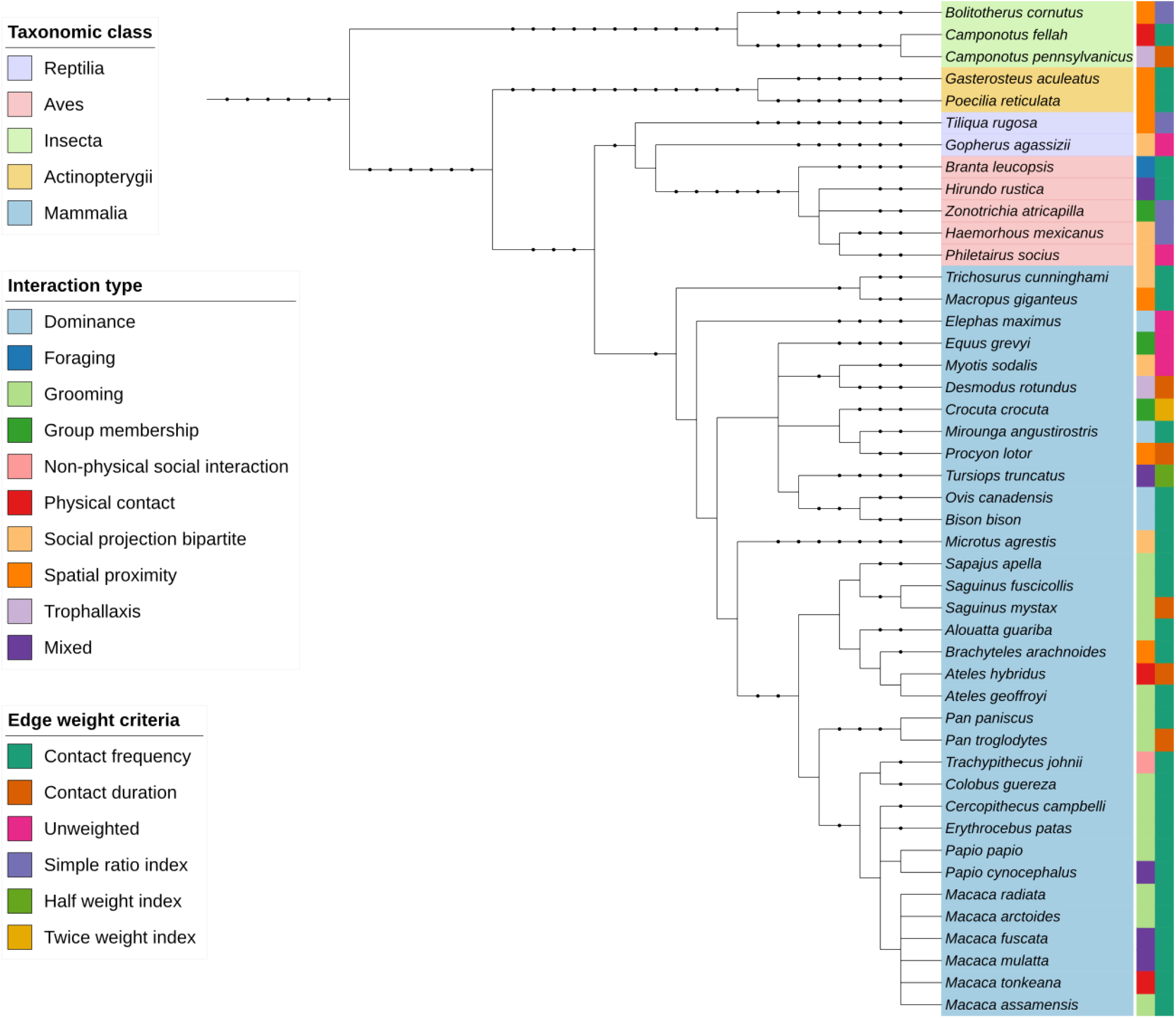
Phylogenetic distribution of non-human species included in the Animal Social Network Repository (ASNR). The first color strip includes the species’ scientific name, and is color coded according to the taxonomic class. The second color strip is coded according to the social interaction quantified in the network, and the third color strip is coded according to the weighting criteria of the network edges.

The geographical locations where data for each social network were collected is shown in Figure 2. The United States contributes the largest number of studies and the repository contains data from Central and South America, Europe, Africa, Asia and Australia. Additionally, most studies are in free-ranging populations.

**Figure 2:**
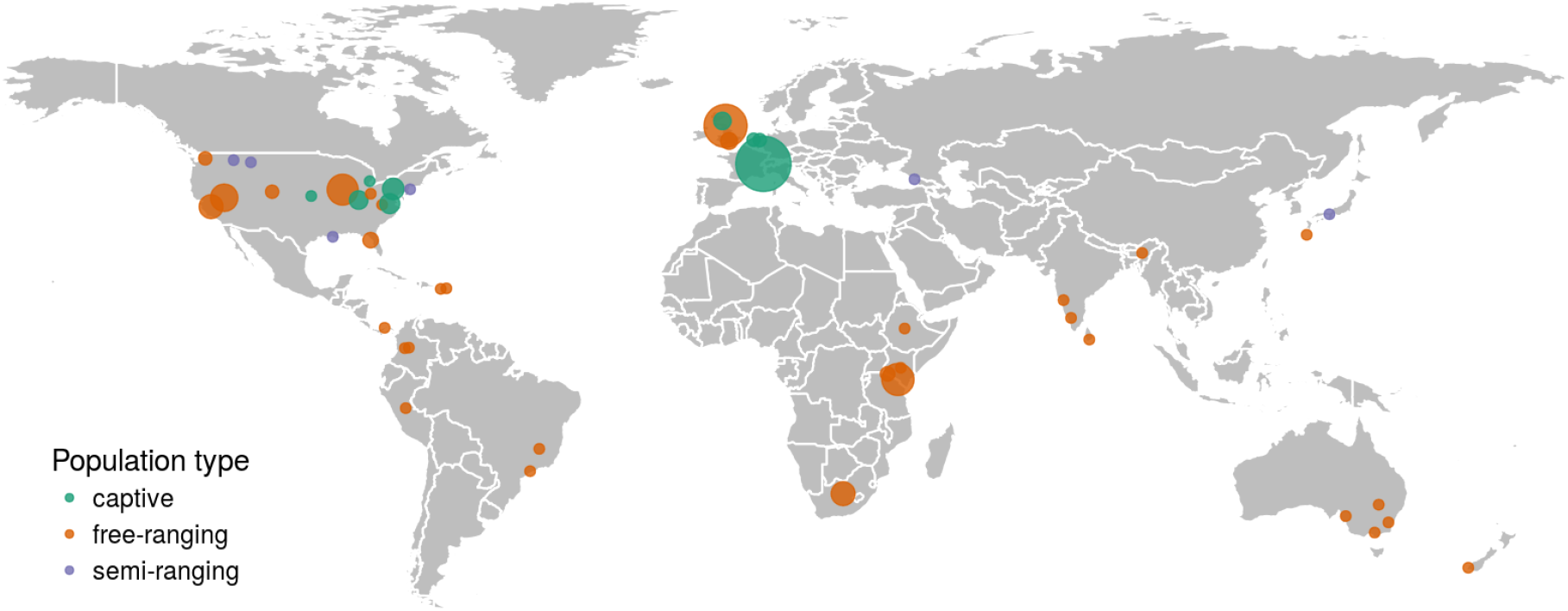
Geographical distribution of the social networks included in ASNR. The points indicate the geographical location where data for each social network was collected. The point size is proportional to the number of social networks collected at each location. Point color denotes whether the monitored animal populations were captive, semi-ranging or free-ranging.

### Behavioral types

The behavioral data span a range of social interaction types from direct physical contacts such as grooming and trophallaxis to indirect interactions such as spatial proximity and association (Figure 1).

Additionally, contact intensity were distributed across six categories – unweighted (i.e., all edges have weight equal to one), contact frequency, contact duration, simple ratio index [10], twice weight index [9], and half weight index [10] (Figure 1).

### Data collection methodology

Figure 3 summarizes the methodology and data collection techniques described in original data sources that were used to collect the networks. We highlight that studies rely on a variety of data collection methodologies and timescales, reflecting empirical constraints and the disparate scientific purposes of each study. It is important that future comparative studies take into account these differences [20].

**Figure 3:**
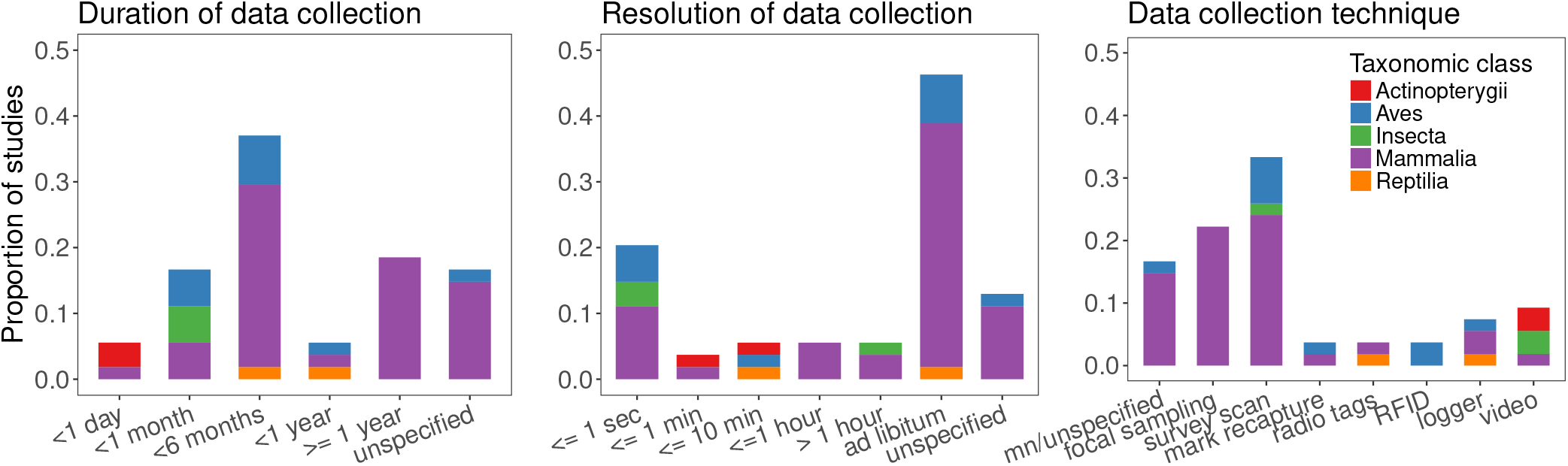
Duration, time resolution and technique of data collection of social networks included in the repository. mn = manual.

### Assessing network structure

We used the Python `NetworkX` package [12] to examine the structural properties of the social networks associated with each species. We calculated the following structural properties for each social network in the repository: total nodes, total edges, network density, network average degree, degree heterogeneity, degree assortativity, average clustering coefficient (unweighted and weighted), transitivity, average betweenness centrality (unweighted and weighted), average clustering coeffcient (weighted and unweighted), Newman modularity, maximum modularity, relative modularity, group cohesion, and network diameter. These network metrics are defined in Table 1.

**Table 1:**
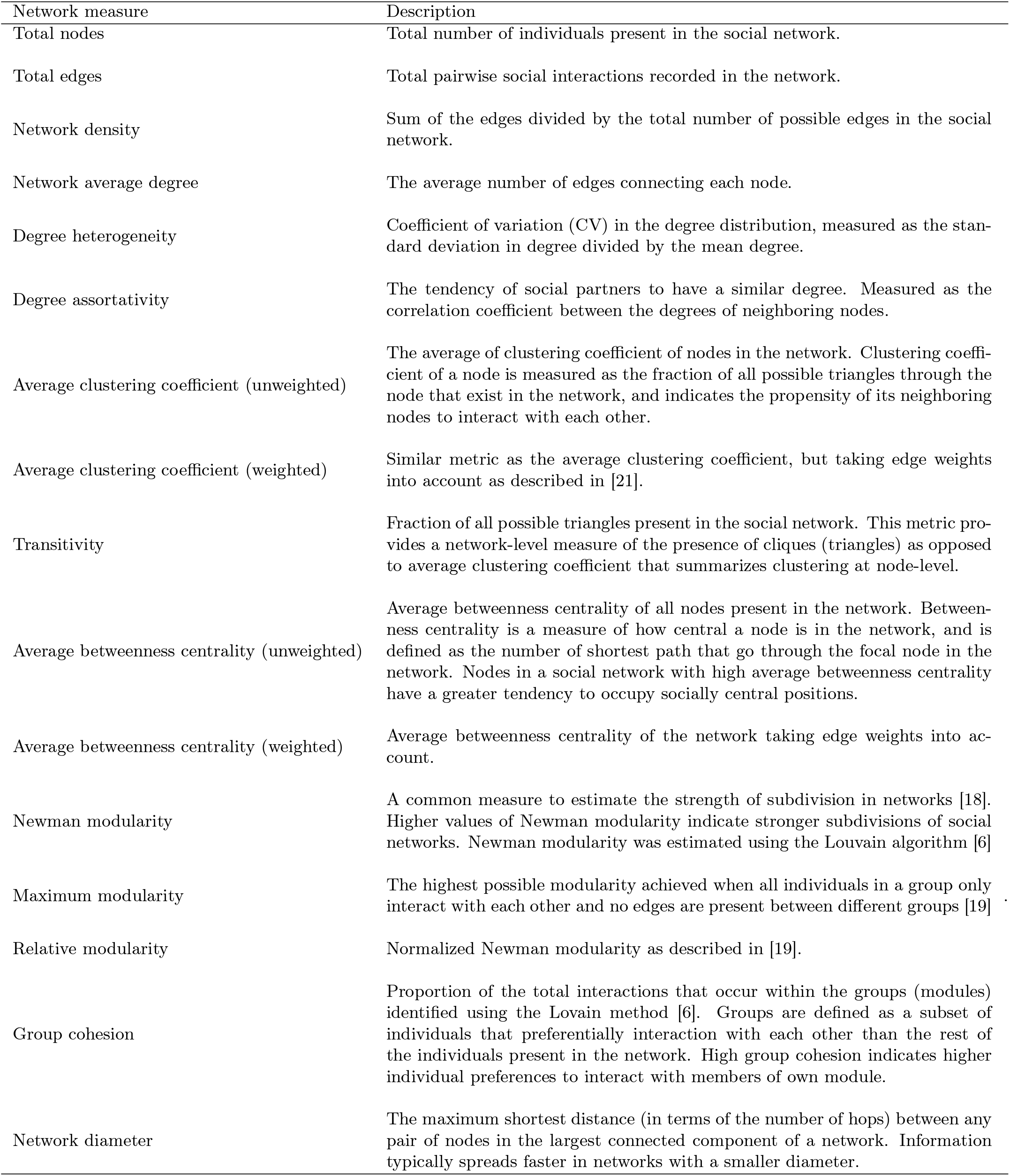
Structural properties of the networks described in ASNR.

In Figure 4 we capture the structural similarity between the social networks included in the repository. Social networks of mammals tend to cluster together, although some structural overlap also exists with the social networks of insects and fish. Social networks that describe spatial proximity, physical contact and grooming interactions between animals tend to be structurally similar.

**Figure 4:**
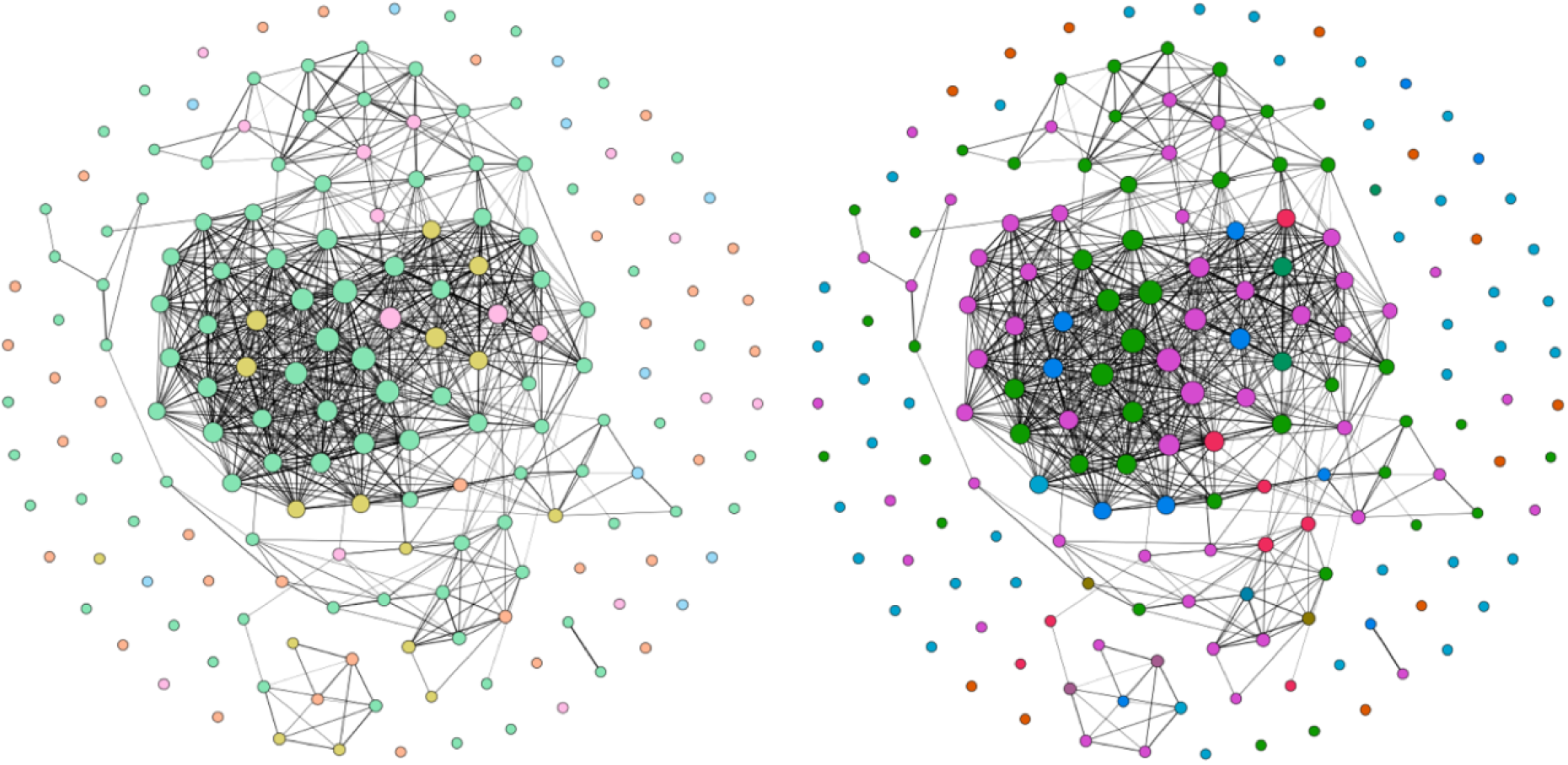
Graphical representation of similarity of networks based on six network metrics – degree heterogeneity, network density, average clustering coefficient, degree assortativity, betweenness centrality and relative modularity. Each node in the network represents a unique social group of an animal species, and an edge between two nodes demonstrates the similarity of their network structure. If a social group contained more than one network (for example, snapshots of a temporal network), an average value was calculated for each network metric. A z-score of each network metric was calculated. Two social groups were considered to be structurally similar (and connected by edges) if they were within one standard deviation of each other in the z-score distribution of all six network metrics. The figures on the left and right are identical except for node colors: (left) node colors indicate taxonomic classes. Green – mammalia, orange - ave, pink - actinopterygii, yellow - insecta and blue - reptilia. (right) Node colors indicate type of interaction represented as edges. Pink -spatial proximity, green - grooming, light blue - social projection bipartite, orange - group membership, dark blue - physical contact, red - dominance interaction, dark green - trophallaxis, brown - foraging, purple - non physical social interaction, teal - overall mix.

## Usage Notes

There are several open-source network libraries that can be used to analyze and visualize the networks provided in `GraphML` format at ASNR. Examples of network analysis and visualization softwares include `NetworkX` in Python, `igraph` in R, `Cytoscape`, `yEd` and `Gephi`.

## Acknowledgements

This work was supported by the National Science Foundation Ecology and Evolution of Infectious Diseases Grant 1216054.

